# Channel expression aligns with dendritic conductance load in the reinforcing, not compensatory, direction in cortical GABAergic neurons

**DOI:** 10.64898/2026.11.11.765852

**Authors:** Giansalvo Cirrincione, Filippo Grassia

## Abstract

Neurons could offset the conductance load imposed by their dendrites through channel expression (compensation), or channel expression could move with that load (reinforcement); which arrangement holds in adult cortex is unknown. Writing rheobase as the product of input conductance and an effective-voltage factor turns the question into a sign test. In mouse cortical GABAergic Patch-seq neurons, after removal of transcriptomic type and covariates, an out-of-fold channel-gene score co-varies positively with residual input conductance (median Spearman *ρ* = 0.32 over 20 cross-fitting splits, *n* = 493) and with a conductance score predicted from dendritic morphology alone (*ρ* = 0.23, *n* = 483). The association is carried mainly by Sst and Pvalb cells and is close to zero in Vip cells. Its slope on the compensation axis is 0.13, against − 1 for compensation and 0 for independence, so the finding concerns direction rather than magnitude: it excludes a population-wide compensation rule without implying strong amplification. Reduced NEURON models show that a compensatory correction is biophysically feasible, and the morphology-derived conductance predictor transfers from mouse to human reconstructions (*ρ* = 0.27, *n* = 128).

**Impact statement:** In adult cortical inhibitory neurons, ion-channel expression varies with dendritic conductance load in the direction that adds to its effect on excitability rather than offsetting it, weakly but consistently.

**Research organism:** Mouse, Human

## 1 Introduction

Neurons with different morphologies and conductance profiles can produce similar electrical outputs. This degeneracy is a central theme in cellular and systems neuroscience, where multiple parameter combinations may yield comparable activity, robustness and homeostatic function [1– 7]. At the same time, dendritic structure imposes biophysical constraints on input resistance and current flow. Classical cable theory predicts that membrane load shapes input resistance, and compartmental modelling has long shown that dendritic structure can influence neuronal output [8–15]. A claim that morphology affects input resistance is therefore not, by itself, a contribution.

Patch-seq data make a sharper question possible. Because morphology, electrophysiology and transcriptomics are observed in the same cells, one can ask how dendritic load and channel expression are *organized* relative to each other once transcriptomic cell-type structure is accounted for [16, 17, 21, 22]. Integrated Patch-seq studies have already shown that morphology, electrophysiology and transcriptome covary across and within transcriptomic types: morpho-electric-transcriptomic types of mouse visual-cortex GABAergic cells [18], continuous phenotypic variation within transcriptomic types of mouse motor cortex [19], and type-specific morpho-electric signatures of human neocortical GABAergic interneurons [20]. These studies establish that the modalities are co-organized; they do not ask in which direction channel expression lies relative to the specific correction that would hold excitability constant against dendritic load. That direction is the object of the present test. The relevant question is directional: is channel expression arranged to *offset* morphology-driven conductance load, stabilizing a common output (homeostatic compensation), or to move *with* it, sharpening type-specific excitability (specialization)?

The question can be made exact. The starting point is the identity

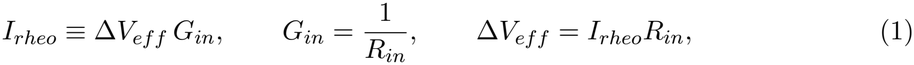

a re-coordinate representation of (*R*_*in*_, *I*_*rheo*_) that defines a conductance branch *G*_*in*_ and an effective-voltage branch Δ*V*_*eff*_ . At constant rheobase,

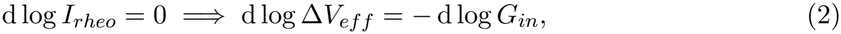

which fixes the direction channel expression would have to take to compensate a morphology-driven conductance perturbation. The test is cross-modal: the perturbation is estimated from SWC geometry, and the potential corrective factor is sought in independently measured channelgene expression. Alignment in the sign of Eq. (2) indicates compensation; the opposite sign indicates reinforcement.

The results are organized around one claim and its controls. After transcriptomic type and covariates are removed, a residual conductance branch organized by dendritic morphology persists; broad channel-class averages do not project onto it, whereas an out-of-fold sparse channel-gene score does, and it does so with the *positive* sign, opposite to the compensatory direction. The sign is reproduced when measured conductance is replaced by a morphology-only prediction, and it is positive in each transcriptomic subclass (significantly so in Sst and Pvalb). The association is small, and the claim concerns its sign, not its size. Reduced NEURON models show that the compensatory correction is biophysically feasible, so the observed sign is not explained by an impossibility of compensation in model neurons, and the morphology-derived conductance predictor transfers from mouse to human reconstructions.

The conclusion is therefore directional and limited: in this adult cross-sectional Patch-seq population, channel expression lies on the side of the conductance branch that adds to dendritic load rather than offsetting it, which is the pattern expected from type-specific specialization and not from a population-wide compensation rule.

## 2 Results

### 2.1 Dataset, analysis counts and branch variables

The primary mouse analysis used matched morpho-electric-transcriptomic Patch-seq data from cortical GABAergic neurons. Different analyses used different subsets because each stage required a different intersection of morphology, electrophysiology, transcriptomics, graph recon-structions or NEURON feasibility. All counts used in the present draft are stated in Table 1. The effective-voltage factor Δ*V*_*eff*_ is a passive-coordinate factor derived from measured rheobase and input resistance, and is read as a branch coordinate rather than as the true biophysical spike threshold.

**Table 1:**
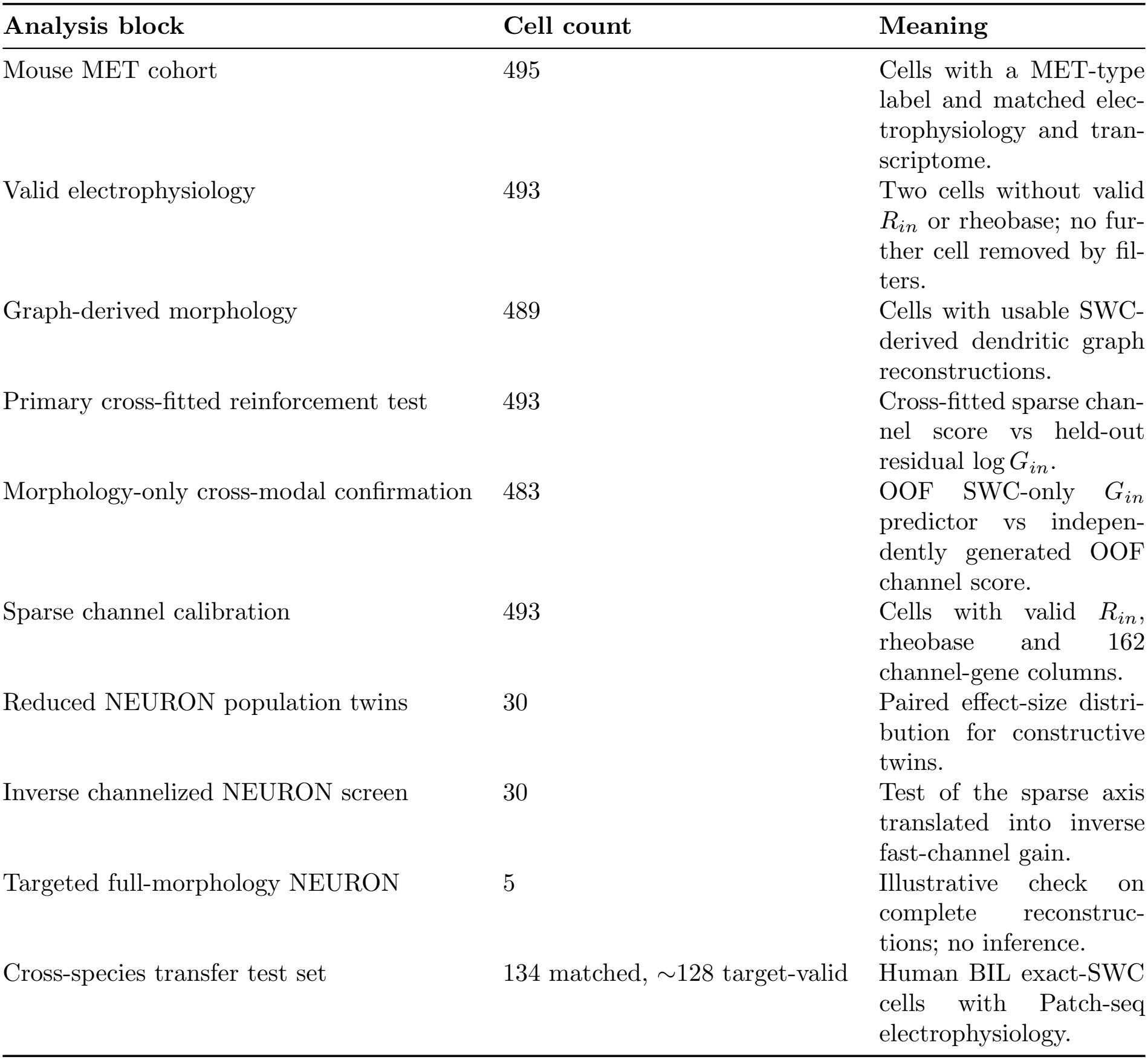
Analysis-specific sample sizes. Each row reports the cell set used for the corre-sponding analysis.

| **Analysis block** | **Cell count** | **Meaning** |
|---|---|---|
| Mouse MET cohort | 495 | Cells with a MET-type label and matched elec-trophysiology and tran-scriptome. |
| Valid electrophysiology | 493 | Two cells without valid R<sub>in</sub> or rheobase; no fur-ther cell removed by fil-ters. |
| Graph-derived morphology | 489 | Cells with usable SWC-derived dendritic graph reconstructions. |
| Primary cross-fitted reinforcement test | 493 | Cross-fitted sparse chan-nel score vs held-out residual log G<sub>in</sub>. |
| Morphology-only cross-modal confirmation | 483 | OOF SWC-only G<sub>in</sub> predictor vs indepen-dently generated OOF channel score. |
| Sparse channel calibration | 493 | Cells with valid R<sub>in</sub>, rheobase and 162 channel-gene columns. |
| Reduced NEURON population twins | 30 | Paired effect-size distri-bution for constructive twins. |
| Inverse channelized NEURON screen | 30 | Test of the sparse axis translated into inverse fast-channel gain. |
| Targeted full-morphology NEURON | 5 | Illustrative check on complete reconstruc-tions; no inference. |
| Cross-species transfer test set | 134 matched, ∼128 target-valid | Human BIL exact-SWC cells with Patch-seq electrophysiology. |

### 2.2 A residual dendritic load signal remains beyond transcriptomic type

Raw input resistance and rheobase were strongly structured by transcriptomic subclass and channel-program correlates. After residualization by subclass, donor, batch, cortical structure, genotype and other covariates, a dendritic load signal persisted: graph-derived proximal load and morphology PC1 were both associated with lower residual *R*_*in*_ and higher residual rheobase. This is the one part of the branch-separation block that is empirically non-trivial — dendritic geometry remains informative beyond type and covariates, and the residual signal is branch-specific. In the residual target spectrum, a dominant morphology/electrotonic mode exceeded the permutation null whereas the second mode did not, so the conservative statement is a dominant residual mode, not an exact one-dimensional law (Fig. 2).

**Figure 1.**
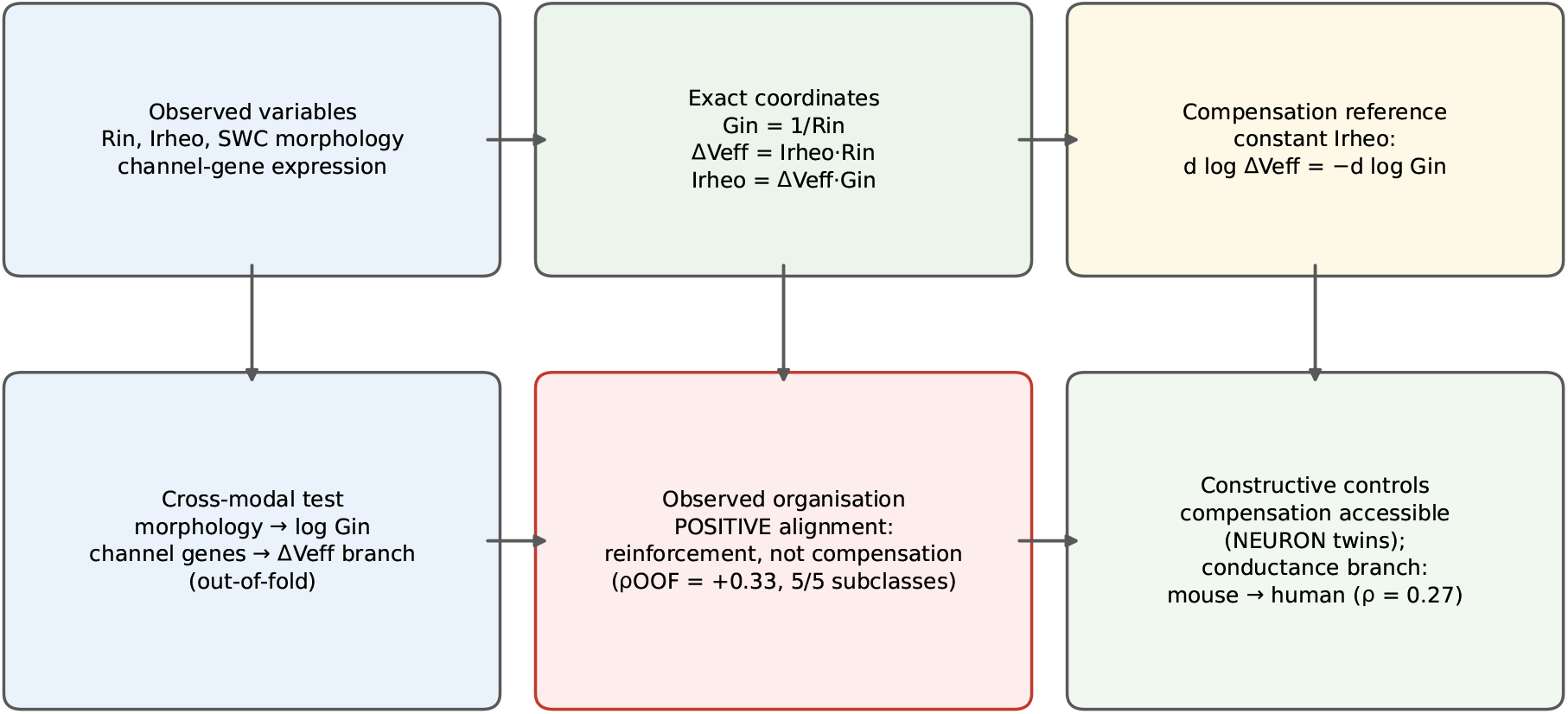
Reinforcement-centered framework. The manuscript is organized around an exact branch identity and a cross-modal sign test. A conductance-branch perturbation is derived from dendritic morphology; a corrective branch is evaluated from independent transcriptomic channel scores; and the sign of their alignment is read against the compensation reference direction. The observed alignment is positive (reinforcement, not compensation), while NEURON twins show that the compensatory direction is biophysically feasible and the conductance branch transfers across species.

**Figure 2.**
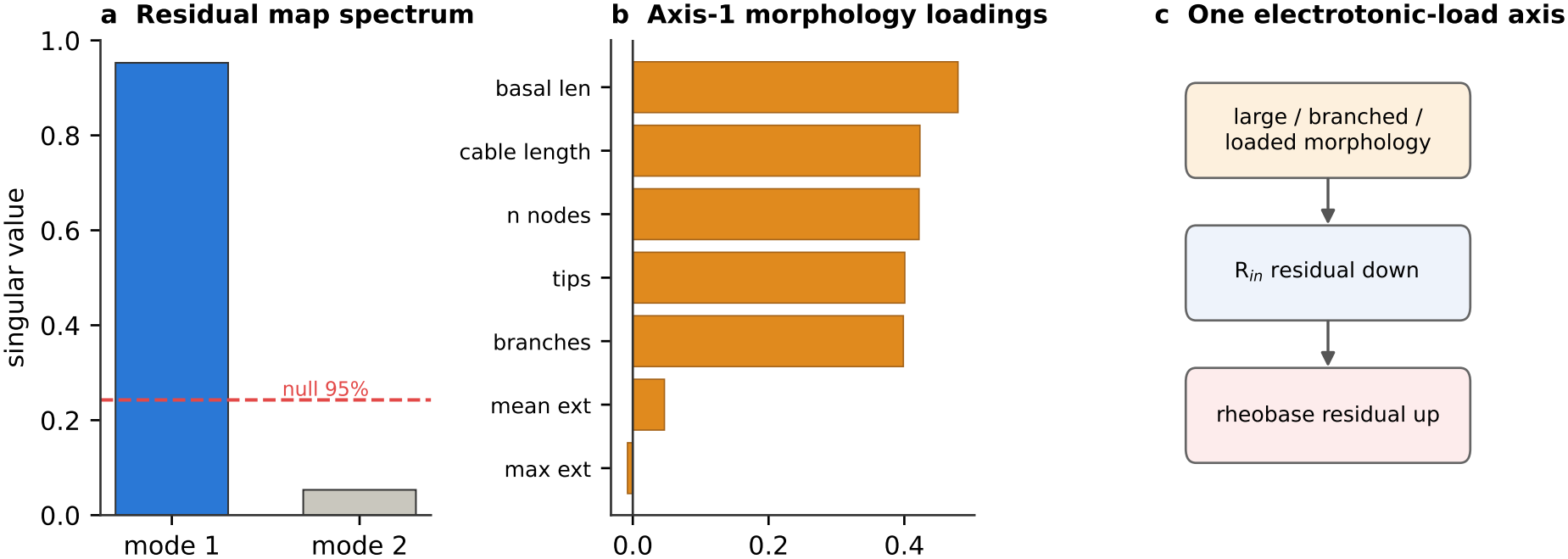
Dominant residual morphology/electrotonic load axis. A residual dendritic-load direction persists beyond transcriptomic type and major covariates. Its direction is cableconsistent, but its role here is as an independently measured, morphology-derived branch for the reinforcement test.

### 2.3 A sparse channel signature co-varies with the conductance branch

A compensation test is only meaningful if channel expression has a functional projection onto the branches at all. Broad class averages did not: a mean fast-spiking channel score did not predict residual Δ*V*_*eff*_ beyond covariates (Spearman *ρ* = 0.017, 95% CI −0.073 to 0.104, *q* = 0.727), and a subthreshold-active average carried only a weak signal (*ρ* = 0.141, *q* = 0.007). A sparse gene-level analysis over all 162 channel genes was therefore performed. Genes and target were residualized beyond subclass and covariates and an elastic-net model was fitted; the final model fitted on all cells selected 37 non-zero genes. In the repeated cross-fitting (Methods), the 100 fold-level models selected a median of 29 genes (range 13–59), and the out-of-fold score carried predictive signal for residual log Δ*V*_*eff*_ (median over splits *R*^2^ = 0.083, range 0.059–0.108; Spearman *ρ* = 0.297, range 0.255–0.322). Selection was reproducible at the level of a core set rather than of the full list: 112 genes entered at least one of the 100 models, 26 entered at least half and 15 at least 80%. All genes selected in at least half of the models kept the same sign whenever selected; only four rarely selected genes (in at most 11% of models) changed sign. The most frequently selected genes were *Kcns3* (100%), *Kcnab3* (100%), *Kcnab1* (98%), *Kcnh7, Scn3b* and *Scn1b* (97%), *Kcnk2* (96%), *Kcnf1* (92%), *Cacna1g* (89%) and *Kcnk3* (88%) (Supplementary file 1).

The out-of-fold channel score is *not* a selective Δ*V*_*eff*_ calibrator. It predicts the residual conductance branch as strongly as the effective-voltage branch it was trained on (median over splits *ρ* = 0.315 with residual log *G*_*in*_ versus *ρ* = 0.297 with residual log Δ*V*_*eff*_ ; *ρ* = 0.380 with residual log *I*_*rheo*_), and every association is *positive* (Fig. 3). The signature is therefore not treated as a conductance-density surrogate or a threshold calibrator. It is a functional transcriptomic axis that co-organizes with both branches, dominated by a conductance-aligned direction. This mixed, positively signed projection is exactly the quantity required by the reinforcement test.

**Figure 3.**
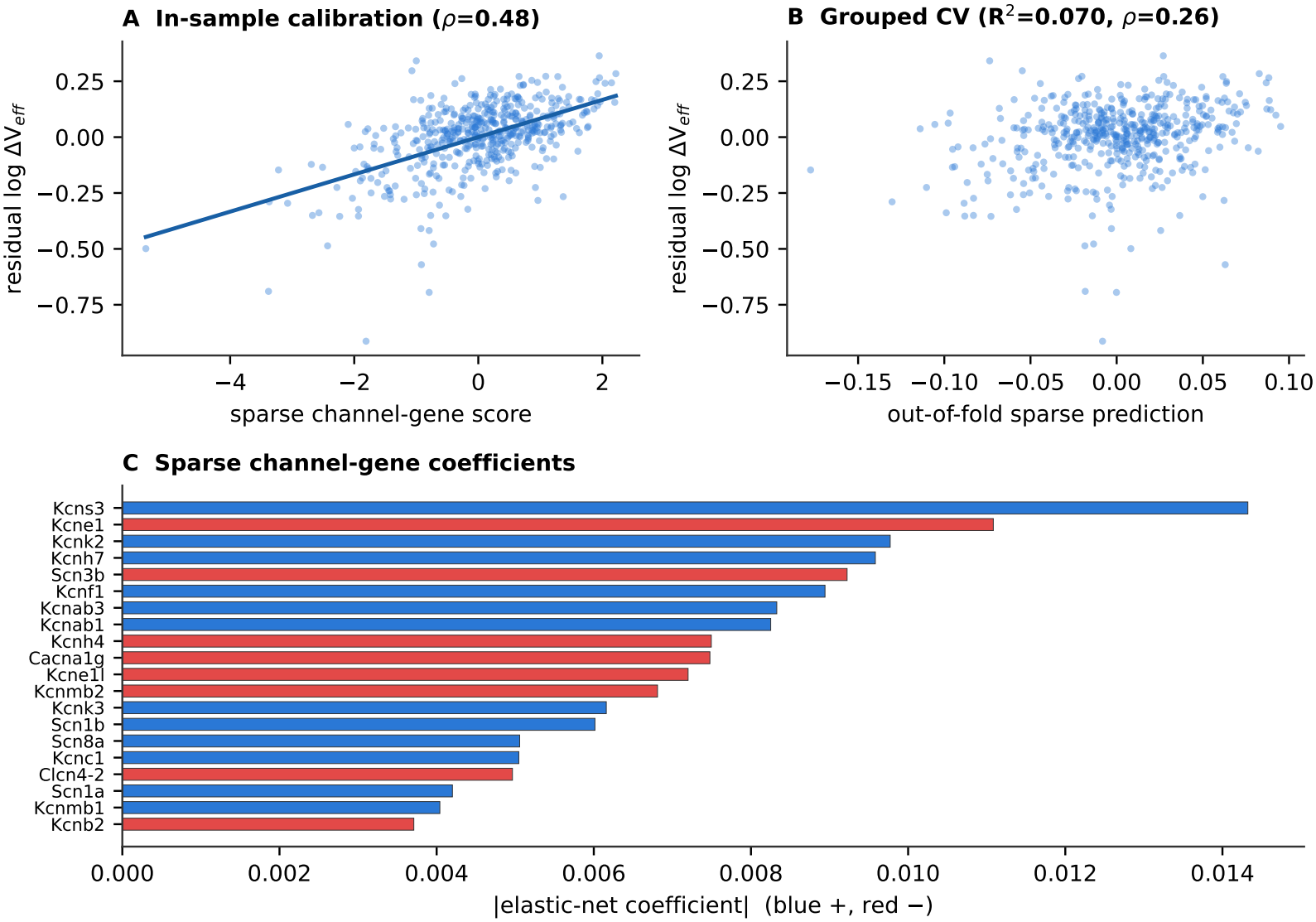
A sparse channel signature co-organizes with the branches. Broad channel-class averages were insufficient, but a sparse channel-gene signature had out-of-fold predictive signal and a monotone residual association. The score is not a direct conductance density and is not a selective effective-voltage calibrator: out of fold it projects onto the residual conductance branch at least as strongly as onto the residual effective-voltage branch, with a shared positive sign. Panel A shows the in-sample calibration curve (*ρ* = 0.48); panel B the grouped cross-validated signal (*R*^2^ = 0.070, *ρ* = 0.26, single cross-fitting split); panel C the most stable selected genes.

To test whether the positive channel-branch alignment requires the specific sparse gene selection or is present in the channel gene space more broadly, a random channel-subset null was computed: 500 times, 29 genes were sampled at random from the 162 channel genes (the median number selected per fold by the real cross-fitted models), and an out-of-fold score was built with the same donor-grouped folds and in-fold target residualization, using ridge regression on the *z*-scored genes in place of the elastic net and without per-gene covariate residualization; the resulting score was tested against residual log *G*_*in*_. The null distribution was positive with median *ρ* = +0.210 and 95th percentile *ρ* = +0.282 (Figure 4—figure supplement 2); the median real cross-fitted association (*ρ* = +0.315) exceeded 99.2% of random subsets. The positive alignment with the conductance branch is thus largely a property of the channel-gene space as a whole: random subsets of the same size already reach about two thirds of the observed association, and the sparse selection adds a modest increment. The sign test below therefore does not depend on the particular genes retained by the elastic net.

**Figure 4.**
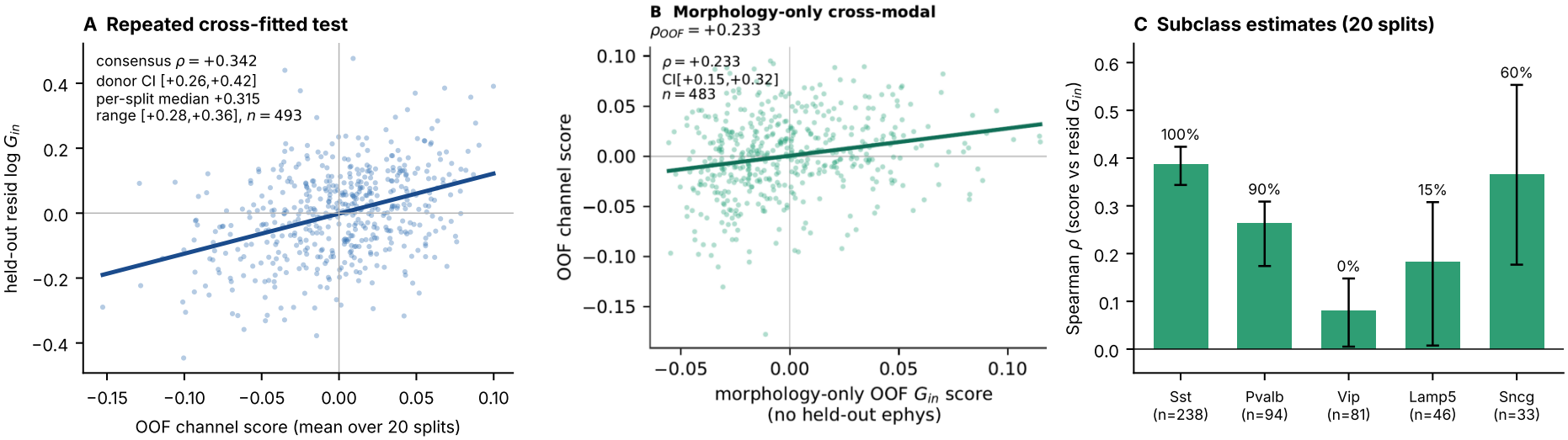
Reinforcing alignment of channel score and conductance branch. A. Repeated cross-fitted measured-conductance test: out-of-fold channel score averaged over 20 splits against held-out residual *G*_*in*_. **B**. Morphology-only test against the single-split channel score. **C**. Subclass medians over 20 splits (bars: 5th–95th percentile; labels: share of splits with *p* < 0.05).

### 2.4 Channel expression aligns with the conductance branch in the reinforcing direction

The compensation condition d log Δ*V*_*eff*_ = − d log *G*_*in*_ fixes a direction. If channel expression compensated morphology-driven conductance load, the channel axis would be arranged *against* the conductance branch — it would carry a negative sign. It does not. Across modalities the sign is positive: the out-of-fold channel axis co-varies with the residual conductance branch with a median Spearman *ρ* = 0.315 over 20 donor-grouped cross-fitting splits (range 0.285–0.355, *n* = 493); the consensus score, averaged over splits, gives *ρ* = 0.342 (*p* < 10^−14^, donor-cluster bootstrap 95% CI 0.261–0.420, 324 donors). This is the opposite of the compensatory prediction. The alignment is not only between-type structure. It persists after removing subclass means (median *ρ* = 0.300, range 0.269–0.339), and the median estimate is positive within each subclass, but with clear heterogeneity (Fig. 4C): Sst +0.386 (*n* = 238, nominally significant in 20/20 splits), Pvalb +0.264 (*n* = 94, 18/20), Sncg +0.365 (*n* = 33, 12/20), Lamp5 +0.183 (*n* = 46, 3/20) and Vip +0.080 (*n* = 81, 0/20). The effect is therefore carried mainly by Sst and Pvalb cells; in Vip cells it is close to zero.

The association is small. A Spearman coefficient of 0.32 corresponds to roughly 10% of rank variance, and 0.23 for the morphology-only test below to roughly 5%. Expressed on the compensation axis, the channel-derived effective-voltage contribution has a slope of +0.13 (median over splits; range +0.10 to +0.16) against the conductance branch, whereas compensation would require −1 and independence 0 (Fig. 4). The slope is numerically much closer to independence than to compensation. What the data exclude is the compensatory direction, together with any sizeable compensatory component: a population-wide compensation rule would place the slope near −1, and the slope is positive in every split. The positive sign is the pattern expected from type-specific specialization; its size does not support a strong amplification of rheobase.

The result is not an algebraic restatement of rheobase variance. In the primary test, the channel axis is an out-of-fold transcriptomic prediction evaluated against held-out residual log *G*_*in*_, whose residual variation is organized by independently measured dendritic morphology. The fully cross-modal confirmation below then replaces measured *G*_*in*_ with a morphology-only held-out predictor, recovering the positive sign in an explicitly SWC-only conductance score. Moreover, a directly measured action-potential threshold barrier — extracted from raw electrophysiological sweeps and independent of the algebraically defined Δ*V*_*eff*_ — follows the same positive sign with the channel axis (*ρ* = 0.089, *p* = 0.049). The effect on this externally measured variable is modest and is reported as directional support, but it argues against a purely tautological interpretation of the reinforcement sign.

To provide a fully cross-modal confirmation, the conductance branch was also evaluated using a morphology-only predictor — without using that cell’s electrophysiological measurements at prediction time (electrophysiology entered only as the training target within the training folds). A ridge-regression model was trained out of fold on compact SWC-derived features (cable length, branch points, tips, node count, Euclidean extent, proximal Sholl mass) to predict residual log *G*_*in*_, yielding morphology-only out-of-fold conductance scores (cross-validated *R*^2^ = 0.093, *ρ* = 0.316 with measured residual log *G*_*in*_; Fig. 4B). This purely morphology-derived score was then correlated with the out-of-fold channel score of the original single cross-fitting split, with neither quantity depending on the held-out cell’s electrophysiology at prediction time. The association was positive (*ρ* = +0.233, *p* = 2 × 10^−7^, 95% CI +0.145 to +0.317, *n* = 483; Fig. 4B), in the reinforcing direction and with a confidence interval that excludes zero. After removing subclass means, the within-population trend remained positive (*ρ* = +0.203, CI +0.116 to +0.292). The morphology-only sign was positive in the three large subclasses (Sst +0.205, *p* = 0.002; Pvalb +0.294, *p* = 0.004; Vip +0.295, *p* = 0.009); the two smallest subclasses (Lamp5 *n* = 45, Sncg *n* = 32) were underpowered for subclass-level inference. The morphology-only cross-modal test therefore replicates the reinforcing direction of the primary result, with no use of the held-out cell’s electrophysiology in the conductance branch.

The positive sign of both tests was retained under donor-level resampling. channel model on the residualized genes of four subclasses and testing on the fifth. The cross-fitted score was positively associated with residual *G*_*in*_ in the held-out subclass for Lamp5 (*ρ* = +0.338, *p* = 0.021), Pvalb (*ρ* = +0.394, *p* = 8.7 × 10^−5^) and Sncg (*ρ* = +0.343, *p* = 0.051). For Vip and Sst the held-out association was weak or negative, consistent with the imbalance induced by withholding large training strata; these results indicate partial, but not complete, leave-one-subclass transfer.

Donor-cluster bootstrap confidence intervals were computed by resampling whole donors (*n* = 493 cells, 324 donors; Figure 4—figure supplement 1). The 95% CI remained positive for the primary measured-conductance test (consensus score, *ρ* = +0.342, CI +0.261–+0.420) and for the morphology-only cross-modal test (*ρ* = +0.182, CI +0.038–+0.338). (*ρ* = +0.325, 95% CI +0.242–+0.407, *p* < 10^−12^) provides the better-powered within-population robustness check.

### 2.5 Compensation is biophysically feasible in model neurons

A positive sign would be uninformative if compensation were physically unavailable. Two checks address this possibility. The required correction lay within one observed channel-score standard deviation for 61.8% of cells, within two for 90.2%, and within the observed 1–99% range for 98.0%: the channel space had ample amplitude to implement compensation, but the population did not occupy that direction.

A population reduced-NEURON screen then converted the compensation rule into constructive counterfactual twins. Real dendritic graphs were reduced to radial cable-like models, a morphology perturbation was applied, and the required Δ*V*_*eff*_ correction was translated into a sodium-conductance correction via the M2 sensitivity d log Δ*V*_*eff*_ /d log *g*_*Na*_ = −0.9167. In 30 cells, morphology-only perturbation produced a median absolute rheobase error of 0.168 log units (95% CI 0.148–0.181); rule-compensated twins reduced this to 0.032 (0.026–0.064, improving 29/30 cells) and mild-rule twins to 0.026 (0.013–0.032), while wrong-sign controls increased it to 0.322 (0.271–0.348) (Table 2). The mild-rule correction, fixed a priori as 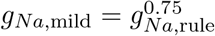, improved 30/30 cells and reached the 20% iso-rheobase tolerance in 30/30 cells; wrong-sign controls were worse than compensated twins in 30/30 cells (paired Wilcoxon, mild-rule versus morphology-only *p* = 8.5 × 10^−7^; versus wrong-sign *p* = 8.7 × 10^−7^). In five complete recon-structions, run as an illustration rather than as a statistical test, the ordering was the same: median absolute error 0.119 after morphology-only perturbation, 0.081 after rule compensation (improvement in 4/5 cells, 2/5 within the 20% tolerance) and 0.280 for the wrong-sign control (worse than the compensated twin in 5/5) (Fig. 5).

**Table 2:** Population reduced-NEURON twin statistics. Absolute log_10_ rheobase errors relative to the baseline twin. Confidence intervals are bootstrap 95% intervals for the median. The wrong-sign control is a negative control.

| **Variant** | **n** | **Median error** | **95% CI** | **Success** < 20% |
|---|---|---|---|---|
| Morphology-only perturbation | 30 | 0.168 | 0.148–0.181 | 0.07 |
| Rule-compensated twin | 30 | 0.032 | 0.026–0.064 | 0.77 |
| Mild-rule compensated twin | 30 | 0.026 | 0.013–0.032 | 1.00 |
| Wrong-sign control | 30 | 0.322 | 0.271–0.348 | – |

**Figure 5.**
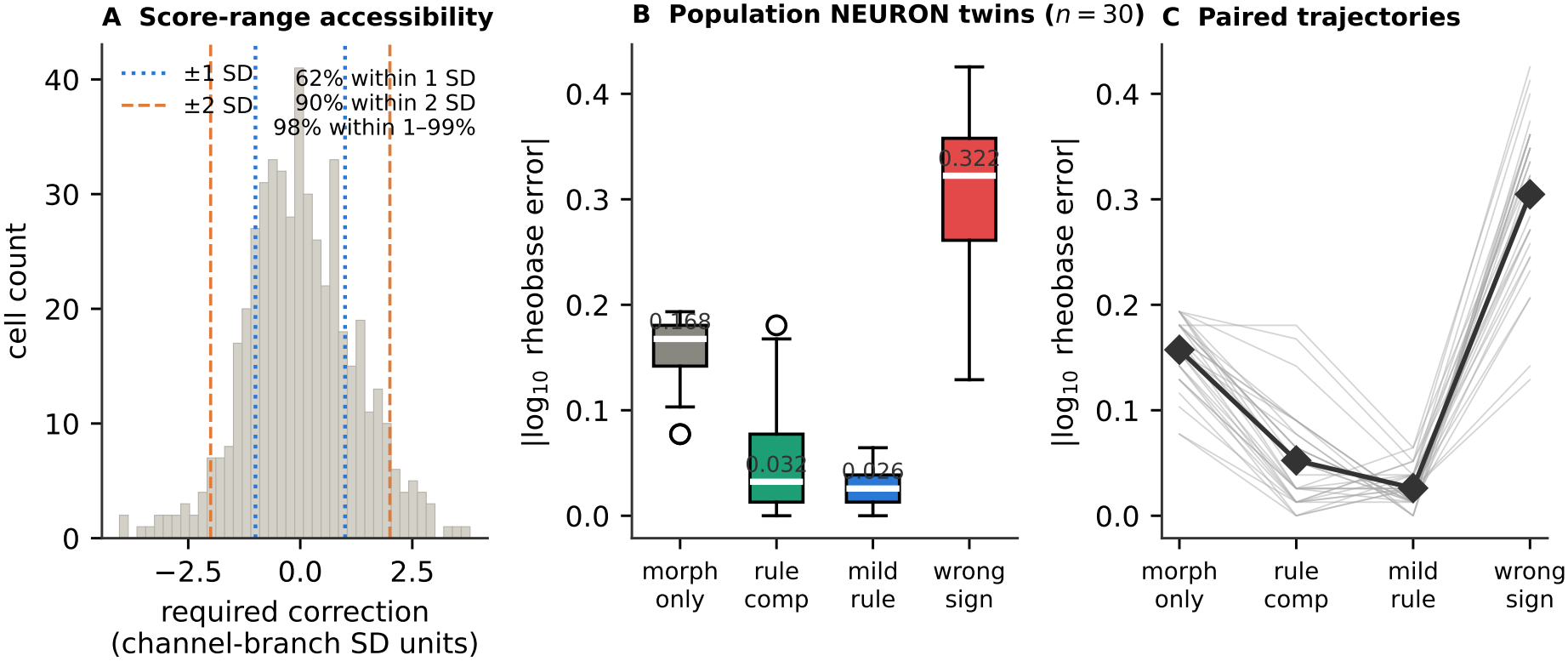
Compensation is biophysically feasible in model neurons. The compensatory correction required to restore iso-rheobase lies within the observed channel-score range for almost all cells (**A**), so the positive sign (Fig. 4) is not explained by an unavailable correction. Reduced NEURON twins built from real dendritic graphs implement the analytic correction: applying the rule moves morphology-perturbed cells back toward baseline rheobase (**B**), whereas wrong-sign controls move them further away.

The scope of this check is limited by its construction. The sodium-conductance correction was derived from the M2 sensitivity, fixed a priori from the same reduced models, so the recovery of rheobase by rule-compensated twins is close to guaranteed once that sensitivity is accurate; the twins demonstrate biophysical feasibility of the correction in model neurons, not that biological neurons could implement it through coordinated changes in channel expression. The informative elements are the wrong-sign controls, which rule out a sign-agnostic effect, and the capacity analysis, which shows that the required correction lies within the observed channel-score range. Together they exclude the explanation that the positive sign arises because compensation is unavailable in principle.

### 2.6 A transcriptomic channel axis generates a functional Δ*V*_*eff*_ branch in NEURON

The sparse channel-gene axis was tested against an explicit biophysical readout. The score was translated into an inverse fast-channel gain in reduced NEURON models, 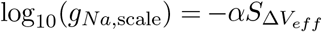, with *α* ∈ *{*0.05, 0.10, 0.15, 0.20*}* fixed a priori and *α* = 0.10 primary. As expected from the imposed monotone mapping, the induced change in simulated Δ*V*_*eff*_ was strongly ordered by the score (*ρ* = 0.988 at *α* = 0.10). The informative test was branch selectivity: although channelization necessarily changed model conductance, the final channelized *G*_*in*_ was only weakly and non-significantly associated with the score (*ρ* = 0.319, *p* = 0.086). The sparse axis thus generated a strong functional Δ*V*_*eff*_ branch while leaving the final conductance coordinate weakly constrained (Fig. 6), a constructive bridge from the transcriptomic signature to the biophysical branch used above.

**Figure 6.**
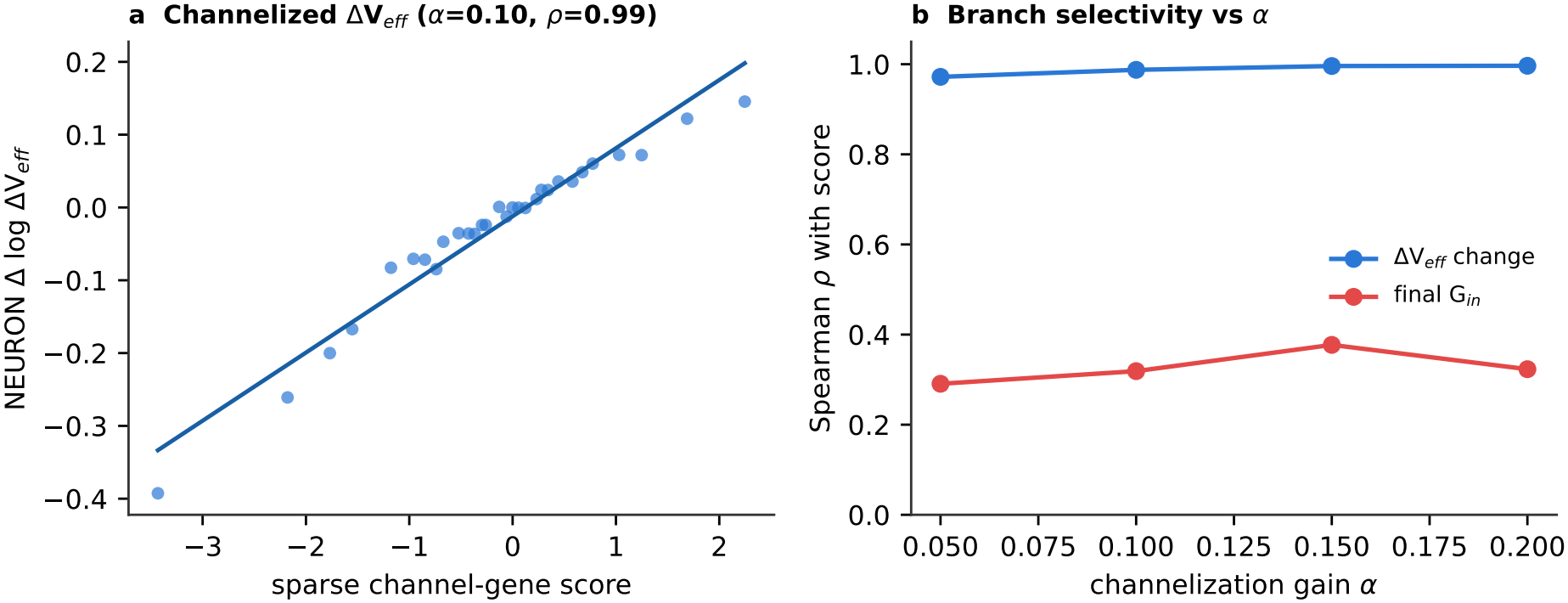
Sparse channel score generates a functional Δ*V*_*eff*_ branch in NEURON. Translating the sparse transcriptomic channel axis into inverse fast-channel gain produces a monotone simulated Δ*V*_*eff*_ response, closing the functional proxy loop without claiming that mRNA is a direct conductance-density measurement.

### 2.7 The conductance branch that anchors the alignment is conserved across species

The reinforcing alignment is anchored by the morphology-derived conductance branch. If this branch reflects a conserved cable-geometric constraint rather than a mouse-specific idiosyncrasy, a predictor trained on mouse morphology should transfer to human as a quantitative, out-of-sample statistic. A regularized predictor was trained to map compact dendritic-load features (cable length, branch points, tips, node count, somatic Euclidean extent and proximal Sholl mass) onto the residual conductance branch using mouse cells only, with features *z*-scored within species (internal cross-validation *R*^2^ = 0.091, Spearman *ρ* = 0.316). The model was then frozen and applied, without any exposure to human labels, to *n* = 128 human Brain Image Library exact-SWC reconstructions.

The frozen mouse predictor transferred. Its output was positively associated with the matched human residual conductance branch (−residual log *R*_*in*_) at Spearman *ρ* = 0.266 (*p* = 0.0024, 95% CI 0.102 to 0.419), and with the raw conductance direction at *ρ* = 0.563 (*p* = 5 × 10^−12^, 95% CI 0.422 to 0.678). A permuted-label null gave |*ρ*|_95%_ ≤ 0.174, which the observed residual transfer exceeds (Fig. 7). This is a quantitative cross-species transfer of the conductance branch, established out of sample. It is not a mouse-to-human test of the reinforcing alignment itself: paired human channel-gene expression is not available in these reconstructions, so the transferred quantity is the conductance branch that anchors the axis.

**Figure 7.**
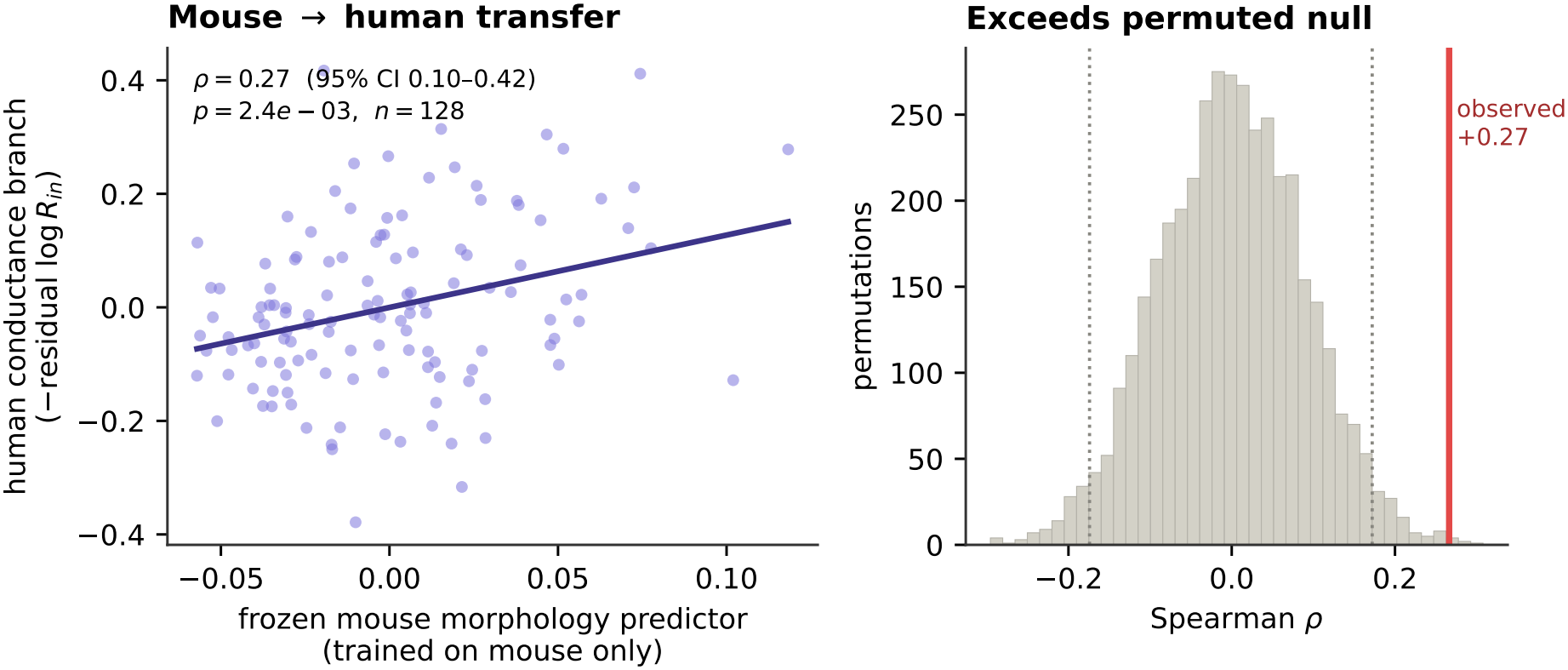
Cross-species transfer of the conductance branch. A morphology predictor trained only on mouse and frozen transfers to human exact-SWC reconstructions (*ρ* = 0.27, 95% CI 0.10–0.42, *n* = 128), exceeding a permuted-label null.

### 2.8 Dendritic operator geometry is a supporting analysis

Dendritic operator geometry was constructed by treating each SWC reconstruction as a rooted metric graph and deriving cable-like operator coordinates, including effective resistance proxies, Green/resolvent access, spectral proximal mass and participation ratios. These features were specific to the conductance branch and did not improve prediction of Δ*V*_*eff*_ ; their incremental value beyond compact linear morphology was small. Operator geometry is therefore not a third central claim. Its role is limited to supporting the interpretation of dendritic load as a conductance-branch coordinate derived from rooted graph structure.

## 3 Discussion

### 3.1 The result is a reinforcing sign

The branch identity *I*_*rheo*_ ≡ Δ*V*_*eff*_ *G*_*in*_ is a coordinate choice, and the association between passive dendritic load and input conductance is expected from cable theory. Neither is the contribution here. The contribution is directional and positive. After transcriptomic type and covariates are removed, a residual dendritic conductance signal persists and defines a morphology-derived conductance branch. Tested against this branch, the channel program does not oppose conductance load, as homeostatic compensation would require; it co-varies with it. Out of fold the channel axis is positively associated with the residual conductance branch (median *ρ* = 0.32 over 20 cross-fitting splits), the association survives within-subclass adjustment, and its median estimate is positive in each subclass, strongest in Sst and Pvalb and near zero in Vip. The adult cross-sectional population therefore lies on the reinforcing side of the compensation reference: channel expression co-varies with dendritic load in the direction that adds to, rather than offsets, its effect on rheobase. This is the pattern expected from type-specific specialization rather than from a common-output stabilization program, and it is the central claim of the manuscript.

The claim rests on the sign, not on the magnitude. The out-of-fold associations explain about 5–10% of rank variance, and the slope on the compensation axis (+0.13) is far closer to independence than to compensation. A residual signal of this size is expected after transcriptomic type and the main covariates are removed, but it means that most of the cell-to-cell variation in the effective-voltage branch is not organized by dendritic load in either direction. The defensible reading is a weak but consistent reinforcing trend that excludes a population-wide compensation rule; it is not evidence that channel programs strongly amplify morphological differences.

### 3.2 Why the reinforcement result is not a circular restatement of rheobase

Because Δ*V*_*eff*_ and *G*_*in*_ are coordinates of (*R*_*in*_, *I*_*rheo*_), a regression of one algebraic component on the other would be tautological. The reinforcement test is not that regression. The conductance branch is measured through one modality — dendritic morphology — and the corrective branch through another — out-of-fold channel-gene expression. The compensation law fixes the sign in which these independent modalities would have to align if the population stabilized output. They align in the opposite sign. This cross-modal construction is what separates a compensation test from a variance decomposition of rheobase. The test is further anchored by a directly measured action-potential threshold barrier, independent of the algebraic Δ*V*_*eff*_, which follows the same positive sign with the channel axis (*ρ* = 0.089, *p* = 0.049). The effect is modest and reported as directional support, but it argues against a purely circular interpretation: the reinforcement sign appears on a quantity that is not a re-expression of the identity.

### 3.3 The positive sign is not forced by a biophysical constraint

The positive sign is not explained by an inability to compensate. Two lines of evidence indicate that compensation is physically available. First, the correction required to restore iso-rheobase lies within the observed channel-score dynamic range for almost all cells. Second, reduced NEU-RON twins built from real dendritic graphs implement the analytic correction and construct counterfactual iso-rheobase cells, whereas wrong-sign controls move rheobase further away. Because the correction in the twins was derived from a sensitivity fixed a priori in the same models, this establishes feasibility in model neurons rather than a biological mechanism. Within that scope, the population does not occupy the compensatory direction even though it is within reach of the observed channel space.

This is compatible with, and complementary to, the degeneracy and homeostasis literature [2, 4, 5, 7]. The present analysis addresses a narrower question: whether an adult cross-sectional Patch-seq snapshot is arranged along a global morphology–channel compensation manifold. It is not; channel expression lies on the reinforcing side. This says nothing about compensation on developmental or activity-dependent time scales, in other cell states, under perturbation, or at the level of local protein trafficking and channel density. The two notions also refer to different objects. Homeostatic compensation in the sense of these studies is a dynamic process within a cell, acting over hours to days after a perturbation, whereas the present test reads a static population covariance across cells at one adult time point. A cell can compensate dynamically around its own set point while the set points of different cells are arranged in a reinforcing pattern; the two are not mutually exclusive. The present result constrains the second, not the first: it shows that the adult population is not arranged along the cross-cell compensation direction, while leaving within-cell homeostatic regulation open.

### 3.4 Channel mRNA is a functional co-organizing axis, not a conductance surrogate

The reinforcement test does not require, and the manuscript does not assert, that mRNA equals conductance density. The broad class-average analysis shows why such an equation would be too crude: a fast-spiking class average does not project onto either branch. Only a sparse gene-level signature does. That signature is deliberately not treated as a selective effective-voltage calibrator, because out of fold it projects onto the conductance branch at least as strongly as onto the effective-voltage branch and tracks residual rheobase most strongly of all. It is therefore a functional transcriptomic axis that co-organizes with both branches, dominated by a conductance-aligned, reinforcing direction. Two independent controls prevent this from becoming a proxy artifact: the signature has genuine out-of-fold predictive signal, and when translated into an inverse fast-channel gain in NEURON it generates the expected functional Δ*V*_*eff*_ response (*ρ* = 0.988) while leaving the final channelized *G*_*in*_ only weakly constrained (*ρ* = 0.319, *p* = 0.086).

### 3.5 The conductance branch is conserved across species

The reinforcing alignment is anchored by the morphology-derived conductance branch, and that anchor is not mouse-specific. A predictor trained on mouse morphology and frozen transfers out of sample to human Brain Image Library exact-SWC reconstructions, predicting the human conductance branch at *ρ* = 0.27 (95% CI 0.10–0.42, *n* = 128) and exceeding a permuted-label null. This is a quantitative cross-species transfer of the conductance branch only; it indicates that the morphology-to-conductance relation used here reflects a conserved cable-geometric constraint, but it does not test the channel side of the alignment in human cells. A stronger future test would transfer the channel-derived corrective branch once matched human transcriptomic channel data are available.

### 3.6 Limitations

Several limitations bound the claim. First, the effect sizes are small (Spearman coefficients between 0.23 and 0.32, compensation-axis slope +0.13) and support a weak directional trend rather than a strong or one-dimensional law; the effect is carried mainly by Sst and Pvalb cells and is near zero in Vip cells. The headline estimate also depends on how cells are split into folds (range 0.29–0.36 over 20 splits), which is why medians over repeated splits are reported. Second, the population NEURON statistic uses reduced radial cable-like models with a correction derived from a sensitivity fixed in the same models, and the five full-morphology cells are illustrative only; the twins establish feasibility, not a biological mechanism. Third, the channel signature is transcriptomic and sparse: inverse channelization demonstrates functional activity in NEURON but does not identify local protein density, and the externally measured threshold barrier provides only modest directional confirmation. Fourth, cross-species transfer is established for the conductance branch and for *n* = 128 human cells, not for the channel side of the alignment. Fifth, dendritic operator geometry is retained as a supporting interpretation, not as an independent evidentiary pillar. Sixth, the study was not preregistered. The reinforcement test was formulated after exploratory analyses of the same public cohort, and several secondary analyses are reported (subclass estimates, donor-cluster and leave-one-subclass-out resampling, a random channel-subset null, the threshold barrier and the morphology-only test); none is corrected for multiplicity, and the threshold result (*p* = 0.049) would not survive such a correction. Table 3 separates out-of-fold, cross-modal statistics from in-sample and model-based quantities. Replication of the sign test in an independent Patch-seq cohort is the decisive next step. These limitations shape the claim: the manuscript does not assert that biological neurons cannot compensate morphology through channels, but that in adult cortical GABAergic Patch-seq neurons channel expression co-varies with the morphology-derived conductance branch in the reinforcing direction, weakly but consistently, even though the compensatory direction is feasible in model neurons.

**Table 3:** Locked numerical values. OOF, out-of-fold or cross-modal; IS, in-sample; M, model-based; CF, cross-fitted; LOSO, leave-one-subclass-out.

| **Analysis** | **Kind** | **n** | *ρ* / **R**<sup>2</sup> | **95% CI** | **Notes** |
|---|---|---|---|---|---|
| Primary, median of 20 splits | OOF | 493 | +0.315 | range [0.285, 0.355] |  |
| Primary, consensus score | OOF | 493 | +0.342 | [+0.261, +0.420] | donor bootstrap |
| Morphology-only cross-modal | OOF | 483 | +0.233 | [+0.145, +0.317] | single split |
| Within-subclass, median of splits | OOF | 493 | +0.300 | range [0.269, 0.339] |  |
| Compensation-axis slope, median | OOF | 493 | +0.13 | range [0.10, 0.16] | vs −1 / 0 |
| Donor-cluster bootstrap (morph-only) | OOF | 483 | +0.182 | [+0.038, +0.338] | single split |
| Random channel-subset null | OOF | 500 | 99.2% | null med. +0.210 | 29 genes |
| Score vs Δ*V*<sub>*eff*</sub>, median | OOF | 493 | +0.297 | range [0.255, 0.322] | *R*<sup>2</sup> = 0.083 |
| Threshold barrier | OOF | — | +0.089 |  | single split |
| Sparse signature in-sample *ρ* | IS | 493 | +0.482 | [+0.408, +0.551] | calibration |
| Morphology-only internal CV | OOF | 483 | *R*<sup>2</sup> = 0.093 |  | SWC features |
| Mouse→human transfer | OOF | 128 | +0.266 | [+0.102, +0.419] | *p* = 0.002 |
| LOSO, held-out subclass | OOF | — | 3/5 |  | Sst, Vip ≈ 0 |

*Subclass medians (splits with p* < 0.05*): Sst* +0.39 *(20/20), Pvalb* +0.26 *(18/20), Sncg* +0.37 *(12/20), Lamp5* +0.18 *(3/20), Vip* +0.08 *(0/20). Genes in* ≥ 80% *of 100 models: 15.*
*NEURON twins (M, n* = 30*): morph-only 0.168, rule 0.032 (29/30), mild-rule 0.026 (30/30), wrong-sign 0.322; M2 sensitivity* −0.9167*. Full morphology (M, n* = 5*): 0.119, 0.081, 0.280.*
*Channelization (M): ρ* = 0.988 *at α* = 0.10*; final G<sub>in</sub>: ρ* = 0.319*, p* = 0.086*.*

### 3.7 Conclusion

An exact branch coordinate system was used to turn a qualitative question about morphology– channel organization into a directional, falsifiable test. The contribution is not the algebraic identity, nor the cable-theoretic fact that dendritic load lowers input resistance. It is that channel expression is organized in the reinforcing, not the compensatory, direction: out of fold, over repeated cross-fitting splits and with positive median estimates in every transcriptomic subclass, the sparse channel branch aligns *positively* with the morphology-derived conductance branch, also when that branch is predicted from morphology alone. The alignment is weak, and its slope on the compensation axis is close to independence; what it excludes is a population-wide compensation rule. A directly measured action-potential threshold barrier follows the same sign, with marginal support. Reduced NEURON twins show that compensation is biophysically feasible in model neurons, and a mouse-trained, frozen morphology predictor transfers out of sample to human exact-SWC reconstructions, indicating that the conductance branch anchoring the alignment is conserved. Replication of the sign test in an independent cohort, and a human test of the channel side, are the natural next steps.

## 4 Materials and methods

### 4.1 Mouse Patch-seq data and preprocessing

All primary sparse-channel statistics reported in the Abstract, Results and Section 4.10 derive from the fully cross-fitted pipeline described in Section 4.3; the morphology-only cross-modal confirmation is described in Section 4.8.

The mouse data are the Allen Institute Patch-seq recordings of GABAergic neurons in mouse visual cortex [18]: electrophysiology from the DANDI Archive (DANDI:000020), transcriptomes from the Neuroscience Multi-omic Archive (counts-per-million release of 13 May 2020) and cell metadata from the Allen Institute. The 495 cells with a morpho-electric-transcriptomic type label were assembled into a matched table with cell identifiers, transcriptomic labels, electro-physiology, dendritic morphology and channel-gene expression. Input resistance was restricted to physiologically plausible values (20 *< R*_*in*_ *<* 1000 MΩ) and rheobase required positive. The measured variables were *R*_*in*_ and *I*_*rheo*_; the derived variables *G*_*in*_ = 1*/R*_*in*_ and Δ*V*_*eff*_ = *I*_*rheo*_*R*_*in*_. Because *G*_*in*_ = 1*/R*_*in*_, residual log *G*_*in*_ = −residual log *R*_*in*_ under linear residualization, which defines the matched human target in the transfer test. Covariate residualization used transcriptomic subclass, donor, batch, cortical structure, genotype, dendrite type and biological sex when present, encoded as one-hot design matrices with ridge regression (penalty selected by internal cross-validation over 10^−4^ to 10^4^). Cells with a missing value in any covariate, in *R*_*in*_ or in rheobase were excluded rather than imputed; a missing channel-gene expression value was set to zero before the log_2_(1 + *x*) transform, consistent with the count nature of the data.

### 4.2 Graph-derived morphology and dendritic operator coordinates

SWC morphologies were represented as rooted graphs with segment coordinates, radii and edges. Graph-derived proximal load used proximal Sholl-like mass, branch points and terminal counts. Operator geometry was constructed by radial contraction of the rooted graph and extraction of cable-like invariants (effective resistance proxies, Green-function access, proximal spectral mass, shell entropy, participation ratios), used only to support the conductance branch.

### 4.3 Channel partitions and the sparse channel-gene signature

Channel genes were grouped by prefix into passive leak-like (Kir, K2P, HCN, Cl-related), subthreshold-active (KCNQ/Kv7, KCNA/Kv1, KCND/Kv4 and auxiliaries), and fast-spiking (sodium, fast potassium, BK/SK, calcium-associated) classes. Broad class averages were computed by log-transforming expression, *z*-scoring genes and averaging within class. Because class averages did not project onto the branches, a sparse gene-level signature was estimated over all 162 channel genes: genes and target were residualized beyond covariates, and residual log Δ*V*_*eff*_ was predicted by elastic-net regression with grouped cross-validation for out-of-fold predictions; stability selection used bootstrap resampling with repeated elastic-net fits. The primary sparse-channel statistics reported in the Abstract, Results and Section 4.10 were obtained from a fully cross-fitted pipeline. In each outer fold, covariate residualization of the target (log Δ*V*_*eff*_ ) and channel-gene features, feature *z*-scoring, hyperparameter selection (elastic-net *ℓ*_1_/ *ℓ*_2_ ratio and regularization strength), feature selection and model fitting were performed using training cells only. The outer loop used five folds grouped by donor (repeated as described below). Within each training fold the elastic net was tuned by four-fold internal cross-validation over *ℓ*_1_ ratios *{*0.10, 0.25, 0.50, 0.75, 0.90, 0.98*}* and 60 regularization strengths log-spaced between 10^−4^ and 10. The frozen residualization, scaling and fitted model were applied unchanged to the held-out cells to generate out-of-fold channel scores. Because the estimates depend on the assignment of donors to folds, the whole procedure was repeated over 20 random donor-grouped five-fold splits. Folds were built by taking donors in a random order and assigning each to the fold with the fewest cells, so that the splits do not depend on library version. For each statistic the median and range over splits are reported. A consensus score, the mean out-of-fold prediction of each cell over the 20 splits, was tested against the mean held-out residual log *G*_*in*_, with 5000 donor-cluster bootstrap resamples for its confidence interval. Gene-selection stability was summarized as the fraction of the 100 fold-level models in which each gene had a non-zero coefficient. These held-out scores were tested against held-out residual log *G*_*in*_ and other branch variables without any additional fitting.

An earlier analysis, in which covariate residualization was performed once on all cells before the cross-validation loop while feature selection, tuning and fitting remained within training folds, gives the same sign and similar magnitudes; it is retained only as a sensitivity check, because global residualization can slightly inflate the cross-validated *R*^2^. The list of most stable genes in the Results comes from the bootstrap stability selection of this earlier analysis. The out-of-fold score was tested, without further fitting, against the residual conductance branch, effective-voltage branch and rheobase, the residual of log *I*_*rheo*_ being the sum of the two branch residuals under the shared linear residualization. Its out-of-fold projection onto residual log *G*_*in*_ is at least as strong as onto residual log Δ*V*_*eff*_, with a shared positive sign.

### 4.4 Reinforcement test against the compensation direction

The identity yields d log *I*_*rheo*_ = d log Δ*V*_*eff*_ +d log *G*_*in*_, so at constant rheobase the compensatory direction is d log Δ*V*_*eff*_ = − d log *G*_*in*_. Cable theory supplies the expected sign of d log *G*_*in*_; it does not predict whether channel-expression variation aligns with − d log *G*_*in*_ across a Patch-seq population. The conductance branch was taken as residual log *G*_*in*_ and the channel branch as the out-of-fold sparse channel score, scaled into log Δ*V*_*eff*_ units by its in-sample calibration slope for display only. Alignment was quantified by the out-of-fold Spearman association (the robust statistic), by its sign within each subclass and after removing subclass means, and by the ordinary least-squares slope on the compensation axis (−1 compensation, 0 independence, positive reinforcement). For the slope, the out-of-fold score was mapped into log Δ*V*_*eff*_ units by its linear calibration against residual log Δ*V*_*eff*_, and the calibrated contribution was regressed on residual log *G*_*in*_; the 95% interval was reported as the median and range over the 20 cross-fitting splits. As an external check independent of the algebraic Δ*V*_*eff*_, the score was also tested against a directly measured action-potential threshold barrier from raw sweeps.

### 4.5 NEURON compensated twins

NEURON simulations used cable-like reduced models derived from real dendritic graphs and, for targeted confirmation, complete morphology models. Passive properties used *R*_*a*_ = 100 Ω cm and *C*_*m*_ = 1 *µ*F/cm^2^; Hodgkin–Huxley channels used baseline somatic *g*_*Na*_ = 0.12 S/cm^2^ and *g*_*K*_ = 0.036 S/cm^2^. Input resistance was measured from a small hyperpolarizing step and rheobase by bisection of a depolarizing step. After a morphology perturbation changed *G*_*in*_, the required Δ*V*_*eff*_ correction was translated into a sodium-conductance scaling using the M2 sensitivity d log Δ*V*_*eff*_ /d log *g*_*Na*_ = −0.9167. Four variants were compared (morphology-only, rule-compensated, mild-rule, wrong-sign); the mild-rule exponent was fixed a priori at 0.75.

The primary endpoint was absolute log_10_ rheobase error relative to the baseline twin, with a 20% iso-rheobase tolerance for success counts.

### 4.6 Frozen cross-species transfer of the conductance branch

Compact dendritic-load features common to mouse and human exact SWC reconstructions were used: total cable length, branch-point count, tip count, node count, maximum and mean Eu-clidean distance from soma, proximal Sholl crossings at 20–200 *µ*m and Sholl peak crossings. Size features were log(1 + *x*)-transformed and all features *z*-scored within species. A ridge regression (*α* selected over 10^−2^ to 10^3^) was trained on mouse cells only (*n* = 484) to predict residual log *G*_*in*_, with internal *k*-fold cross-validation. The fitted model was frozen and applied, without exposure to human labels, to *n* = 128 human cells; the matched target was −residual log *R*_*in*_. Transfer was quantified by Spearman association with bootstrap 95% confidence intervals and a 4000-draw permuted-label null.

### 4.7 Inverse channelized NEURON calibration

The sparse score was converted into an inverse fast-channel gain, 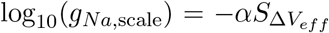, with *α* ∈ {0.05, 0.10, 0.15, 0.20} fixed before analysis and *α* = 0.10 primary. For each cell a baseline reduced model was calibrated, the channelized model simulated, and Δ*V*_*eff*_ recomputed as simulated rheobase times simulated input resistance. The primary statistic was the Spearman association between the score and the induced change in log Δ*V*_*eff*_ .

### 4.8 Morphology-only cross-modal confirmation

To test whether the reinforcement sign could be recovered without using the held-out cell’s measured input resistance, a morphology-only ridge predictor was trained from compact SWC features to predict residual log *G*_*in*_, following the same feature set and scaling as the cross-species transfer (Section 4.6). The analysis was performed in an outer *k*-fold loop. Within each training fold, covariate residualization, within-species feature scaling, ridge hyperparameter selection and model fitting were performed using training cells only; the frozen preprocessing and model were then applied to held-out cells to obtain morphology-only conductance scores. These morphology-only out-of-fold scores were correlated with independently generated out-of-fold sparse channel scores (Section 4.3). For each held-out cell, the two quantities entering the cross-modal test were therefore derived from morphology and gene expression respectively, with no use of that cell’s electrophysiology at prediction time (electrophysiology served as the training target within training folds only). Internal cross-validation quality: *R*^2^ = 0.093, *ρ* = 0.316 between morphology-only predictions and measured residual log *G*_*in*_, *n* = 483.

### 4.9 Statistical reporting and confidence intervals

For the sparse signature, out-of-fold predictions guard against in-sample artifact and the fitted score is reported only as a calibration curve. For the reinforcement test, the reported statistic is the out-of-fold cross-modal Spearman association, its sign within and across subclasses, and a bootstrap interval for the compensation-axis slope. For the NEURON twins, paired within-cell comparisons were emphasized with bootstrap intervals for medians and paired Wilcoxon tests. For the transfer, the model was frozen on mouse before any human evaluation, with significance assessed against a permuted-label null in addition to the bootstrap interval.

### 4.10 Locked numerical values

Table 3 reports all headline numerical values of this version. Values marked “single split” come from the original single cross-fitting split and have not yet been recomputed with the consensus score. The column “Kind” separates statistics computed on held-out cells or across independent modalities (OOF) from in-sample calibration quantities (IS) and quantities obtained from NEURON models (M).

## Data availability

All primary data are public: mouse Patch-seq electrophysiology (DANDI:000020), transcriptomes (Neuroscience Multi-omic Archive, Allen Institute Patch-seq release of 13 May 2020) and metadata from the Allen Institute [18]; human Patch-seq data (DANDI:000636) and human SWC reconstructions (Brain Image Library, DOI:10.35077/g.892). All derived tables, per-figure source data and the outputs of the repeated cross-fitting are deposited at Zenodo (**10.5281/zenodo.24131443**).

## Code availability

All analysis code (Python notebooks that rebuild the matched table from the public sources, run the repeated cross-fitted reinforcement test, the null and leave-one-subclass-out controls, the reduced-NEURON twins, the channelization screen and the cross-species transfer), with a requirements.txt file, random seeds and execution order, is deposited at Zenodo (**10.5281/zenodo.24131443**).

## Author contributions

Giansalvo Cirrincione: Conceptualization, Methodology, Software, Formal analysis, Writing – original draft. Filippo Grassia: Conceptualization, Methodology, Validation, Writing – review and editing.

