## Supplementary_Information for "Channel expression aligns with dendritic conductance load in the reinforcing, not compensatory, direction in cortical GABAergic neurons"

### Figure supplements and Supplementary file 1

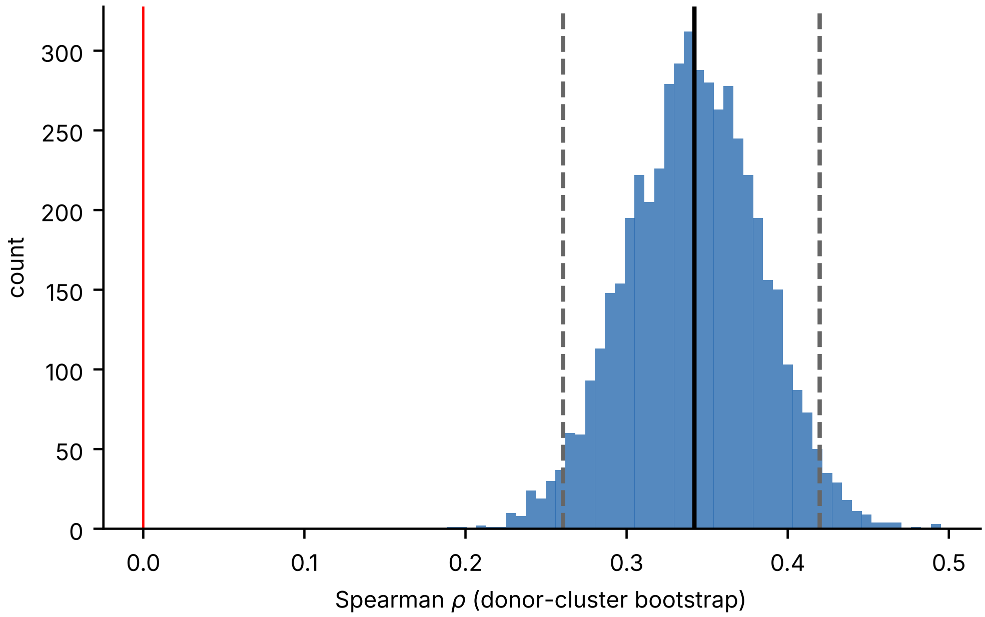


**Figure 4—figure supplement 1. Donor-cluster bootstrap for the primary test.** Consensus out-of-fold channel score (mean over 20 cross-fitting splits) against mean held-out residual $\log G_{in}$; 5000 resamples of whole donors ($n = 493$ cells, 324 donors). Observed $\rho = +0.342$, 95% CI $[+0.261, +0.420]$.

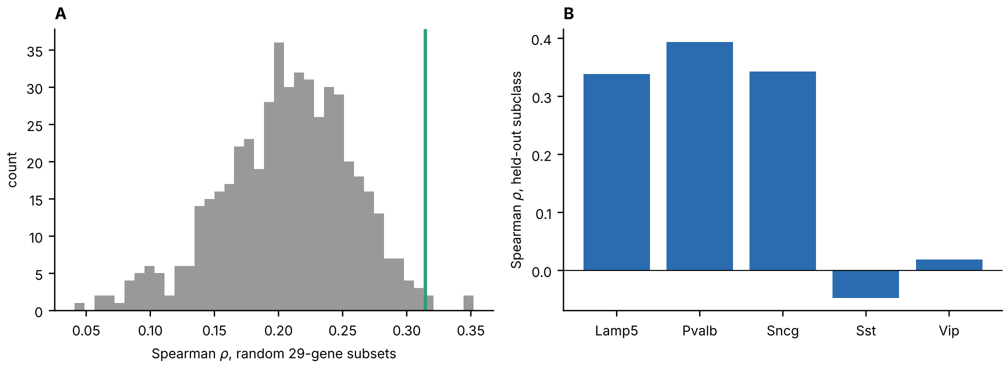


**Figure 4—figure supplement 2. Random channel-subset null and leave-one-subclass-out transfer. A.** Spearman $\rho$ between out-of-fold scores built from 500 random 29-gene channel subsets and held-out residual $\log G_{in}$ (ridge regression, same donor-grouped folds). Null median $\rho = +0.210$, 95th percentile $+0.282$; the median real cross-fitted association ($\rho = +0.315$, green line) exceeds 99.2% of the null. **B.** Leave-one-subclass-out transfer (ridge model on residualized genes): positive in Lamp5, Pvalb and Sncg; close to zero in Sst and Vip.

**Supplementary file 1. Gene-selection stability.** Channel genes selected in at least half of the 100 fold-level elastic-net models (20 cross-fitting splits × 5 folds), with selection frequency and sign of the mean coefficient; every gene kept the same sign in all models in which it was selected.

| Gene | Selected (%) | Sign |
|---|---|---|
| *Kcns3* | 100 | + |
| *Kcnab3* | 100 | + |
| *Kcnab1* | 98 | + |
| *Kcnh7* | 97 | + |
| *Scn3b* | 97 | − |
| *Scn1b* | 97 | + |
| *Kcnk2* | 96 | + |
| *Kcnf1* | 92 | + |
| *Cacna1g* | 89 | − |
| *Kcnk3* | 88 | + |
| *Kcnh4* | 83 | − |
| *Kcnmb2* | 83 | − |
| *Kcnmb1* | 82 | + |
| *Scn1a* | 80 | + |
| *Kcne1* | 80 | − |
| *Kcnc1* | 79 | + |
| *Kcne1l* | 78 | − |
| *Kcnb2* | 77 | − |
| *Scn8a* | 77 | + |
| *Hcn1* | 76 | + |
| *Clcn4-2* | 68 | − |
| *Cacna1b* | 63 | − |
| *Kcna10* | 62 | − |
| *Kcnip1* | 59 | + |
| *Cacnb4* | 56 | + |
| *Kcnj3* | 52 | + |
